# Neither kin selection nor familiarity explain affiliative biases towards maternal siblings in wild mountain gorillas

**DOI:** 10.1101/2022.06.14.496042

**Authors:** Nicholas M. Grebe, Jean Paul Hirwa, Tara S. Stoinski, Linda Vigilant, Stacy Rosenbaum

## Abstract

Evolutionary theories predict that sibling relationships will reflect a complex balance of cooperative and competitive dynamics. In most mammals, dispersal and death patterns mean that sibling relationships occur in a relatively narrow window during development, and/or only with same-sex individuals. Besides humans, one notable exception are mountain gorillas, in which non-sex biased dispersal, relatively stable group composition, and the long reproductive tenures of alpha males mean that animals routinely reside with both same and opposite-sex, and full and half siblings, throughout their lives. Using nearly 40,000 hours of observation data collected over 14 years on 699 sibling and 1258 non-sibling pairs of wild mountain gorillas, we demonstrate that individuals have strong affiliative preferences for full and maternal siblings over paternal siblings or unrelated animals, consistent with an inability to discriminate paternal kin. Intriguingly, however, aggression data imply the opposite. Aggression rates were statistically indistinguishable among all types of dyads except one: in mixed-sex dyads, non-siblings engaged in substantially more aggression than siblings of any type. This pattern suggests mountain gorillas may be capable of distinguishing paternal kin, but nonetheless choose not to affiliate with them over non-kin. A preference for maternal kin occurs despite gorillas not possessing low male reproductive skew, the key characteristic believed to underlie such biases. These results call into question reasons for strong maternal kin biases when paternal kin are identifiable, familiar, and similarly likely to be long-term social partners, and suggest behavioral mismatches at play during a transitional period in mountain gorilla society.

## Introduction

In humans and non-human animals alike, sibling social relationships are marked by continuous dynamics of conflict and cooperation that begin before birth (Trivers, 1974), and can persist throughout an entire lifespan, with important fitness consequences for the individuals involved (Lu, 2007; Hudson & Trillmich, 2008; Nitsch et al., 2013). While classical frameworks of sibling interactions emphasized competition among brood- or litter-mates for limited parental resources during times of dependency (e.g, Mock & Parker, 1997), subsequent developments across numerous academic disciplines (**demography**: e.g. Sear & Mace, 2008; Nitsch et al., 2013; **sociology**: e.g. Steelman et al., 2002; Lu, 2007; **behavioral ecology**: e.g. Silk, 2002; Hudson & Trillmich, 2008; **developmental psychology**: e.g. Lamb et al., 2014) have explored the full arc of sibling competition and cooperation across the lifespan and demonstrated the complexity and diversity inherent to sibling relationships. In understanding the evolution of human sibling relationships in particular, comparative studies of our primate cousins have identified a number of factors predictive of how siblings cooperate and compete. Inconsistent results within and between species, along with the remarkable flexibility of human social systems, however, limits the translational value of many primate models. Here, we address these gaps by presenting an extensive longitudinal study of wild mountain gorillas (*Gorilla beringei beringei*), an endangered ape whose unique, flexible social structure serves as a valuable comparative model to humans.

Classic models of kin selection predict that the social/mating structure of animal groups creates patterns of relatedness between group members, which then selects for kin recognition mechanisms that manifest in differences in cooperative and/or affiliative behavior (Hamilton, 1964; Grafen, 1990; Mateo, 2015). This straightforward idea has spawned a large body of work on kin discrimination in primates, with notably mixed results. Some studies support the existence of sophisticated kin discrimination (Wu et al., 1980; Smith et al., 2003; Widdig et al., 2002, Pfefferle et al., 2014), while others do not, instead favoring simple familiarity as the primary determinant of interaction patterns (Fredrickson & Sackett, 1984; Schaub, 1996; Erhart et al., 1997; Wikberg et al., 2014; Godoy et al., 2016). Inconsistent evidence has led some to suggest that non-monogamous primates evince matrilineal, but not patrilineal, sibling kin discrimination (Mitani et al., 2000; Chapais, 2001; Langergraber et al., 2007). Yet other perspectives challenge this clean distinction, suggesting that complex interactions between familiarity and kin discrimination structure social bonds across primates (see e.g. Silk, 2002; Widdig et al., 2002; Silk, 2009; Lynch et al., 2017).

As one of the main contributors to familiarity, age differences within sibling and non-sibling dyads might influence social dynamics. On one hand, siblings close in age might be more likely to compete for limited parental resources (Tung et al., 2016; Salmon & Hehman, 2021); on the other hand, as longer-lasting co-residents within the same family environment, they might also be expected to form stronger affiliative bonds than siblings distant in age (though, again, this may not apply equally to maternal and paternal sibships; Widdig et al., 2002). It is unclear to what extent age proximity effects are restricted to genetic relatives. Female rhesus macaques appear to bias affiliation towards similarly-aged peers, even when unrelated to them (Widdig et al., 2001). Among female baboons, even in individuals not related through the matriline, dyadic bond strength weakened with increasing age differences; however, when restricting analyses to females unrelated through both the matriline and patriline, effects of age differences attenuated sharply (Smith et al., 2003; Silk et al., 2006). These results once again imply social familiarity (as indexed by age differences) and kin discrimination are both important for predicting sibling relationship qualities (Godoy et al., 2016), though their additive and/or interactive effects remain poorly defined.

Finally, the sex makeup of the dyad might influence interaction styles due to the differential benefits males and females receive from interactions with brothers, sisters, and unrelated partners (e.g. Lonsdorf et al., 2018). From the perspective of males, especially in species who engage in aggressive intrasexual competition, other males, brothers included, can represent important allies (e.g. Meikle & Vessey, 1981; Goodall, 1986) or rivals (Daly & Wilson, 1988) during status-striving efforts in adulthood. In either case, assessing physical capacities or formidability would aid in these efforts. Accordingly, rough-and-tumble play between males might serve important functions as a rehearsal for intrasexual competition in adulthood (Gray, 2019), suggesting such a behavior should occur most often in male-male relationships—a prediction supported by research on male-dominant primates (e.g., Brown & Dixson, 2000; Maestripieri & Ross, 2004). While male-male interaction patterns might generally differ from those of other sex configurations, these differences may themselves partially depend on kinship: in chimpanzees, some evidence suggests that fraternal relationships among adolescents and adults are more affiliative and cooperative than relationships between unrelated males (e.g. Mitani, 2009; Sandel et al., 2020). From the female perspective, evidence for fraternal influences on fitness outcomes is mixed: some research suggests no effect, except perhaps in the case of maternal death (Engh et al., 2009), while one demographic study of humans reports benefits of older brothers on women’s lifetime fitness (Nitsch et al., 2013). From the perspective of both males and females, sisters may represent important future alloparental helpers, either for the individual themselves (e.g. Hamilton et al., 1982; Gould, 2000; Hobaiter et al., 2014), or the individual’s offspring (e.g. Johnson et al., 1980; Nishida, 1983; but see Silk et al., 2006), and thus cultivating relationships with sisters via affiliative interactions might be beneficial for both males and females. These kinds of sex-biased interactions might additionally depend on age differences between siblings (Lonsdorf et al., 2018). Lastly, for females in particular, sororal relationships may exert important influences on future rank and resource acquisition outcomes (Charpentier et al., 2008; Lea et al., 2014; cf. Engh et al., 2009).

Addressing the issues reviewed above, and understanding the nature and evolution of cooperative social relationships in primates more generally, requires long-term investigations of species that reveal how individuals respond behaviorally to socioecological variation (e.g. Alberts & Altmann, 2012). With this principle in mind, mountain gorillas in particular are a compelling comparative model for the evolution and development of human sibling relationships. Long-term monitoring of wild mountain gorillas by the Dian Fossey Gorilla Fund has revealed social structures marked by extensive diversity in relatedness, age proximity, and sex makeup infrequently observed in other primate groups (Robbins et al., 2009; Roy et al., 2014). Mountain gorillas regularly form multi-female, single-male groups, as well as multi-female, multi-male groups in which multiple males reproduce, though paternity data and unsophisticated paternal kin discrimination mechanisms are consistent with historically high reproductive skew (Bradley et al., 2005; Rosenbaum et al., 2015; Vigilant et al., 2015). As a result of this structure, co-resident offspring have a reasonable chance of being full siblings, paternal half-siblings, maternal half-siblings, or unrelated to one another. Like humans, both males and females, upon reaching maturity, may opt to disperse or remain in their natal groups (Robbins et al., 2009; Stoinski et al., 2009), which permits fraternal, sororal, and mixed-sex relationships that can last for an entire lifespan.

In the present study, we use nearly 40,000 hours of behavioral data spanning 14 years to describe patterns of interactions between siblings and demographically-comparable non-sibling dyads in social groups of wild mountain gorillas. Using extensive maternity and genetic paternity data available for 157 identifiable individuals, we examine whether full siblings, maternal half-siblings, paternal half-siblings, and unrelated co-residents exhibit differing patterns of affiliation (playing, grooming, and time spent in close proximity) and agonism (contact and non-contact aggression) in line with models of kin selection, after adjusting for the potential mediating presence of mothers in these interactions. Evidence of relatively weak kin discrimination among gorilla fathers and offspring (Rosenbaum et al., 2015, though see also Vigilant et al., 2015 for evidence of father-daughter inbreeding avoidance) might suggest correspondingly weak social bias toward paternal half-siblings, along with strong bias toward full siblings and maternal half-siblings; we test this speculation for the first time in this species. We also investigate whether familiarity, as indexed by age differences between social partners, predict patterns of affiliative or agonistic behavior, and whether these patterns differ between kin categories—particularly paternal and maternal kin (as some evidence from cercopithecine monkeys suggests; e.g. Widdig et al., 2002).

Lastly, we explore sex makeup as a third axis of variation potentially relevant for understanding sibling and co-resident social relationships. We test whether male-male, female-female, and mixed-sex sibling relationships are characterized by differing rates and types of social interactions, and whether these sex category differences are restricted to kin. Via these comparisons, we ask whether differences are consistent with the kinds of benefits siblings might be expected to deliver later in life: for example, among males, are fraternal relationships marked by higher rates of playing and fighting, and sororal relationships higher rates of grooming?; among females, are sororal relationships marked by the highest rates of grooming compared to any other dyad configuration?; are affiliative patterns unique to siblings, or are comparable trends found in unrelated dyads?

## Results

### Affiliative Behaviors

In our full sample of 1957 unique dyads spanning 7,858 dyad-years, full siblings (n=43 dyads) played and groomed each other significantly more than did paternal siblings (n=555 dyads) or non-siblings (n=1258 dyads; all comparisons *p* < 0.001; Figure 1A, 1B). Maternal siblings (n=101 dyads) played significantly less than full siblings, but groomed at comparable rates. Age differences (in our sample, mean: 5.90 years; SD: 4.57 years; range: 0 – 23.5 years) interacted with relatedness in predicting grooming (*p* = 0.023), but not play (*p* = 0.076). Play consistently dropped for siblings and non-siblings alike as age differences increased (γ ranging from −0.23 – −0.28, all *p* < 0.001; Figure 2A). By contrast, grooming rates were relatively unrelated to age differences between partners (γ ranging from −0.07 – 0.02, all *p* > 0.05; Figure 2B).

**Figure 1.**
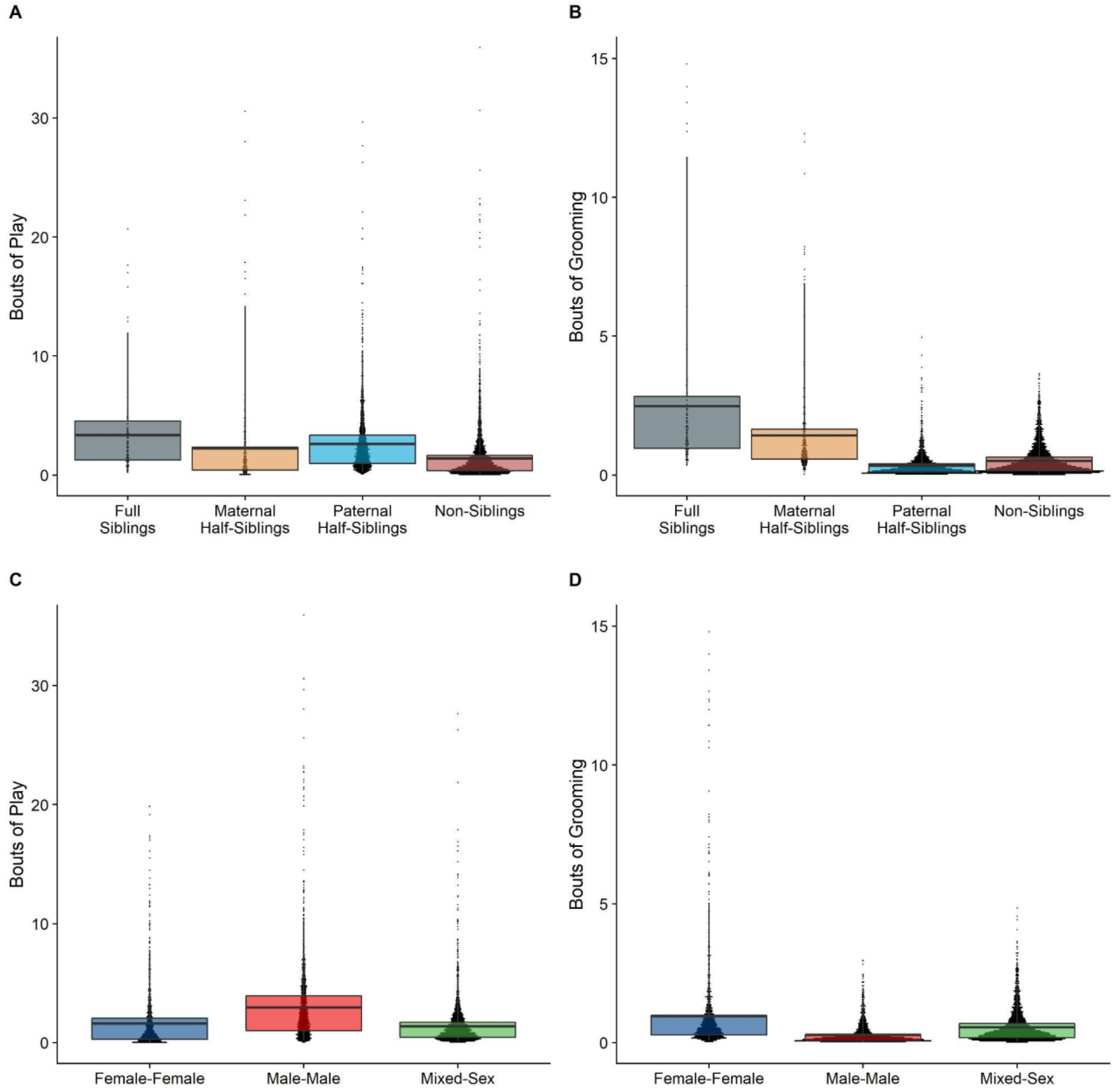
Box and dot plots comparing relatedness categories (A, B) and sex categories (C, D) for play rates (left) and grooming rates (right).

**Figure 2.**
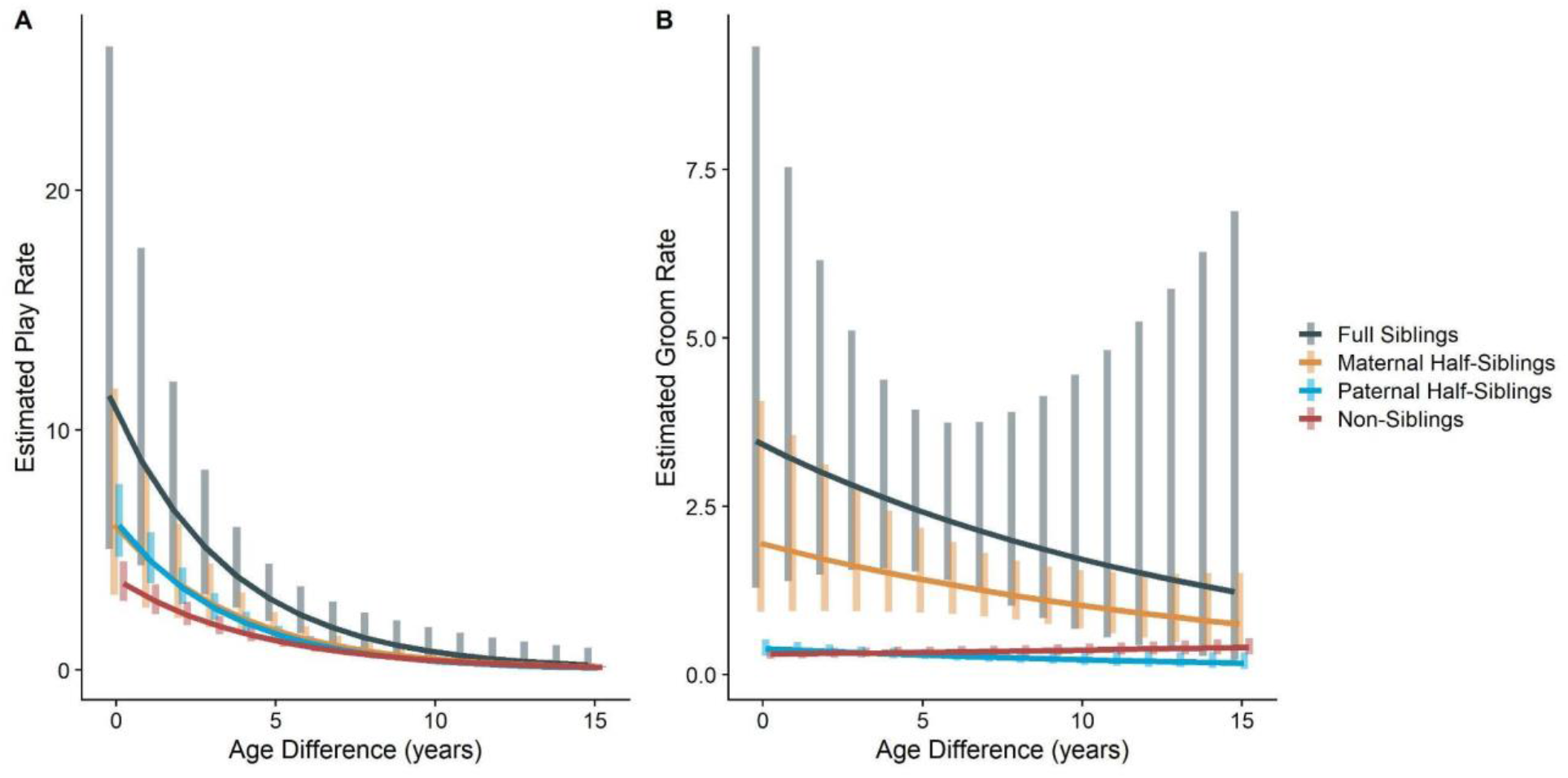
Estimated rates of play (A) and grooming (B) across a range of age differences, separated by relatedness category. Bars represent 95% confidence intervals for rates of behavior at a given age difference.

Male-male dyads (n=520) played more than either mixed-sex (n=981) or female-female dyads (n=456); conversely, female-female dyads groomed each other more than either mixed-sex or male-male dyads (all *p* < 0.001; Figure 1C, 1D). These patterns too were significantly moderated by age differences (*ps* < 0.002), but not relatedness (*p* = 0.078 and 0.112). Play dropped rapidly with increasing age differences (γ = −0.29 – −0.21) for all sex configurations (all *p* < 0.001; Figure 3A). Grooming was steadily low in male-male and mixed-sex dyads (γ = −0.01 and −0.02, *p* > 0.45), though it dropped with increasing age differences in female-female dyads (γ = −0.09, *p* = 0.001), such that differences between sex categories became indistinguishable after approximately a 10-year age difference (Figure 3B).

**Figure 3.**
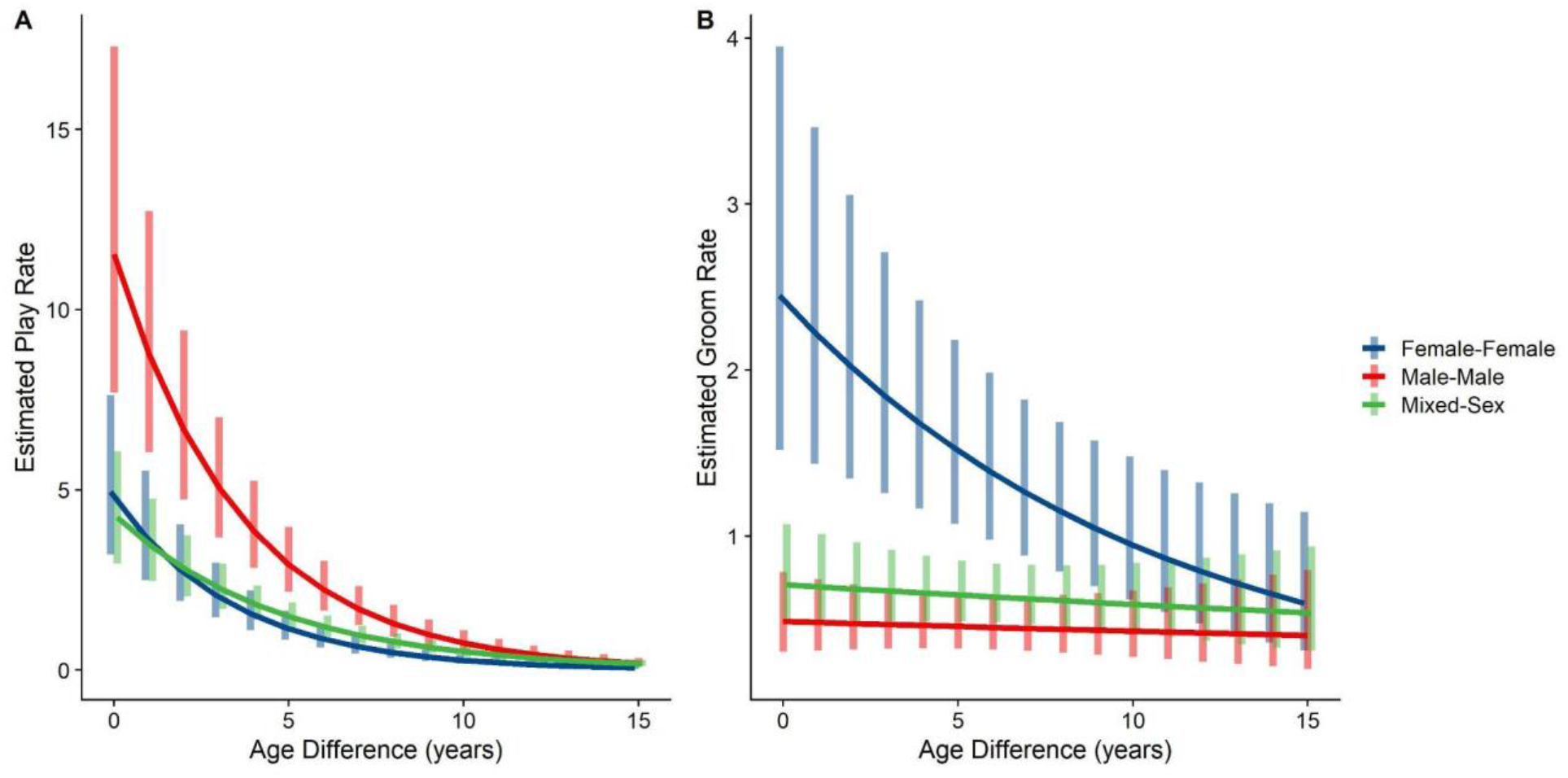
Estimated rates of play (A) and grooming (B) across a range of age differences, separated by sex category. Bars represent 95% confidence intervals for rates of behavior at a given age difference.

### Time spent in proximity

The time dyads spent in close proximity (<2 m) with each other also varied between relatedness categories (*p* < 0.001), with maternal siblings and full siblings once again spending more time near each other than non-siblings, who themselves spent more time in close proximity than paternal siblings did (all comparisons *p* < 0.001; Figure 4A). However, these patterns too were moderated by age differences (*p* < 0.001). Proximity decreased with increasing age differences in maternal siblings and paternal siblings (γ = −0.08 and −0.09,*p* < 0.001), but did not decrease significantly in full siblings or non-siblings (γ = −0.06 and 0.01,*p* > 0.19). Thus, while all classes of siblings spent more time near each other than non-siblings when near in age, even when adjusting for their mother’s presence, this distinction was partially reversed at large age differences, when paternal siblings spent much less time near each other than any other dyad category (Figure 4B).

**Figure 4.**
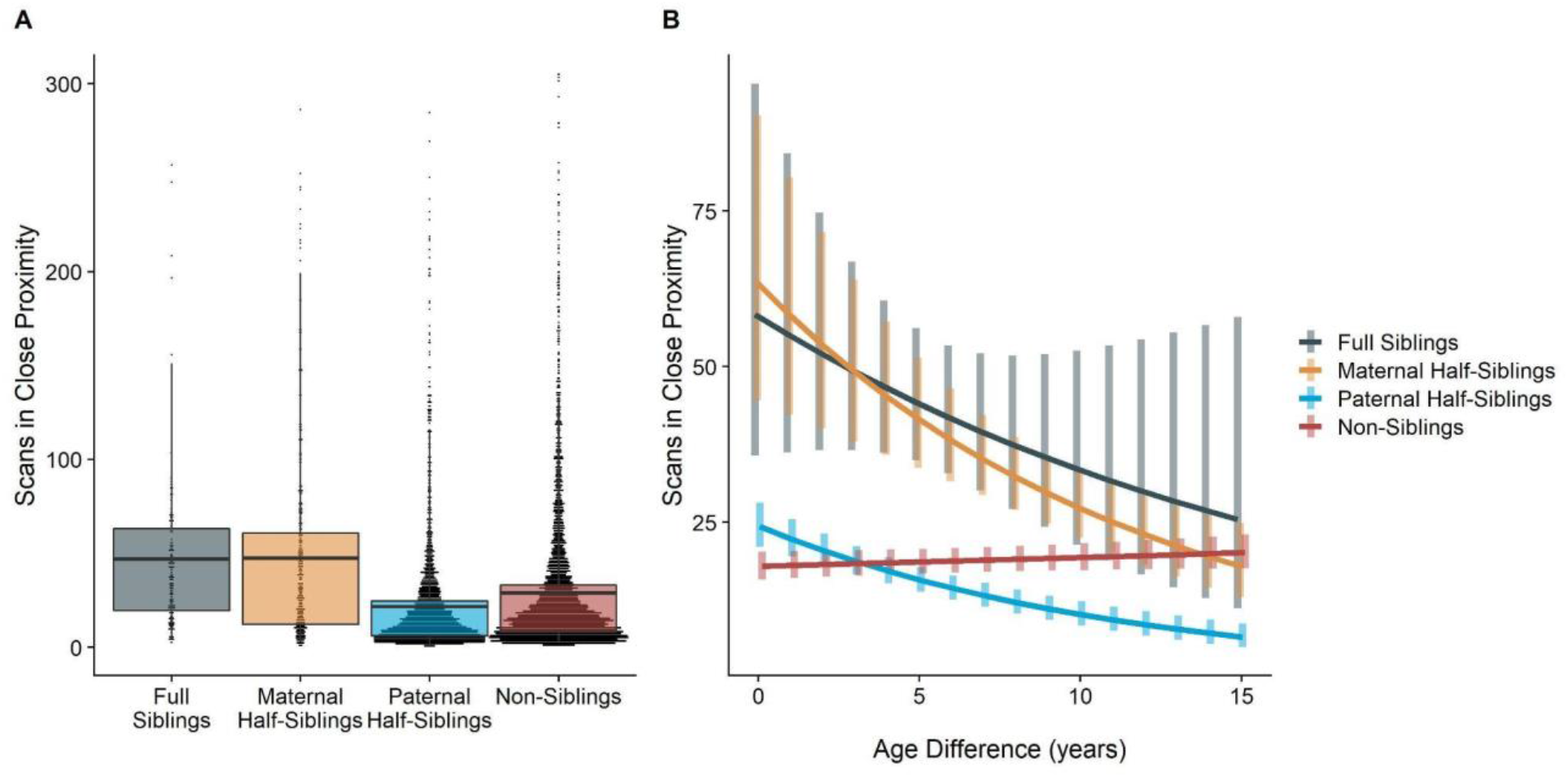
Box and dot plots (A) and estimated trends across a range of age differences (B) for the time gorilla dyads spent in close proximity, separated by relatedness category. Bars in (B) represent 95% confidence intervals for rates of proximity at a given age difference.

### Competitive Behaviors

Neither relatedness nor sex category on their own significantly predicted rates of aggressive behavior (*p* = 0.205 and 0.763, respectively). However, our model did reveal a significant sex makeup × relatedness interaction term (*p* = 0.049; Figure 5A). Decomposing this interaction, among female-female and male-male dyads, there were no statistically significant contrasts between relatedness categories. In mixed-sex dyads, non-siblings engaged in substantially more aggression than any sibling category (all *p* < 0.031).

**Figure 5.**
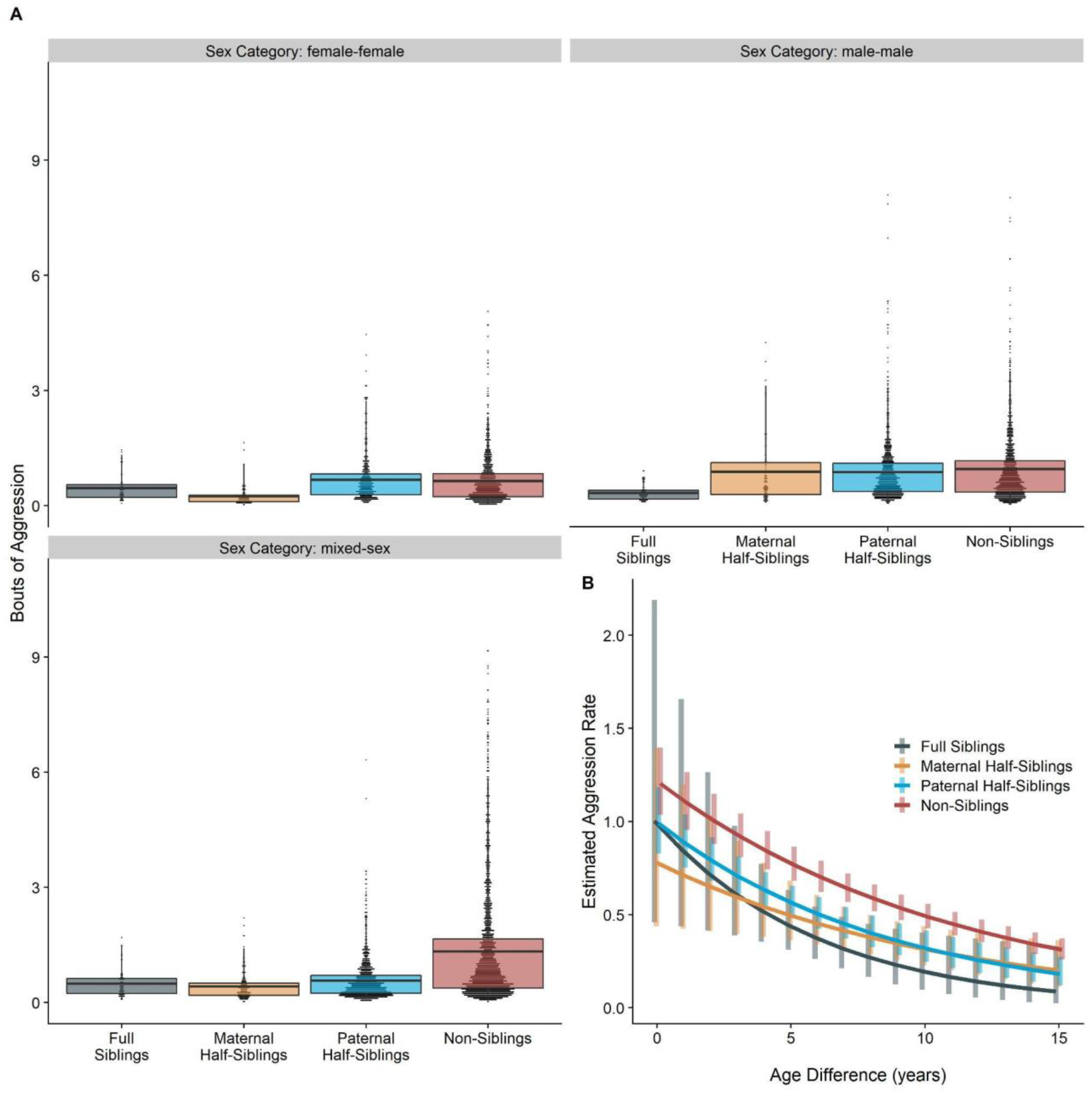
Box and dot plots (A) and estimated trends across a range of age differences (B) for aggression within gorilla dyads, separated by relatedness and sex category. Bars in (B) represent 95% confidence intervals for rates of aggression at a given age difference.

Dyads more distant in age engaged in less aggression than dyads that were closer in age, regardless of relatedness or sex makeup (*z* = −2.11, *p* = 0.035; Figure 5B).

## Discussion

In a comprehensive examination of dyadic mountain gorilla social relationships spanning 14 years and nearly 40,000 hours of observation, we find complex patterns of affiliation and competition within gorilla pairs that speak to sex-, age-, and relatedness-specific social biases. In general, siblings affiliated with each other more and spent more time together than non-siblings, even when accounting for the presence of mothers. But within siblings, affiliative patterns further varied: full and maternal siblings were in most cases much more affiliative than paternal siblings, whose behavior more closely resembled that of non-siblings. We consistently observed a trend for male-male dyads to play more, for female-female dyads to groom more, and for mixed-sex dyads to fall intermediate between these groups. Examining competitive behaviors, on the other hand, revealed a more narrow sibling bias. Aggression was most common in mixed-sex non-sibling dyads, and larger age differences similarly predicted less aggression across all dyad types.

At the broadest level, our results support the existence of affiliative biases towards kin in mountain gorillas. While past research has been largely equivocal about the extent of kin discrimination that relies on mechanisms beyond familiarity (Wikberg et al., 2014; Godoy et al., 2016; Lynch et al., 2017)–and indeed, our results do support a role of familiarity in structuring social interactions–our findings are unlikely to be entirely explained by mere exposure for at least three reasons:

First, gorilla social groups are tight-knit and cohesive compared to their close ape relatives (Goodall, 1986; Remis, 1997; Morrison et al., 2021), such that all individuals in a group, related or not, are very likely quite familiar with one another. Second, our results pertaining to prosocial biases towards siblings are not fully explained by familiarity or exposure, as indexed by age differences (Widdig et al., 2001; Smith et al., 2003; Lynch et al., 2017). We observed clear biases towards kin at all but the largest age differences–and some siblings in our data set were 20 or more years apart in age–even though sibling and non-sibling age-mates in the same social group would typically be expected to possess close familiarity. Finally, while mothers undoubtedly mediate social interactions of offspring, especially for immature individuals, biases towards siblings persist even when adjusting for the frequency of her presence during interaction periods. Jointly, these considerations suggest a sibling bias in mountain gorillas subject to influence, but not determination, by demographic factors, which we interpret as aiding in the development of sibling relationships that exist across timescales rarely observed in other non-human primates.

The observation that full and maternal siblings groomed, played, and spent more time near each other than paternal siblings or non-siblings, who tended to affiliate at comparable rates, further suggests that mountain gorillas, like several other primate species studied (Langergraber et al., 2007; Silk et al., 2006; Lynch et al., 2017), evince much stronger maternal than paternal kin bias (see also Rosenbaum et al., 2015, which found little evidence for paternal kin discrimination among fathers and offspring). Interestingly, this “asymmetric bias” in affiliation seems to persist despite mountain gorillas lacking a key element of the social structure hypothesized to underlie it in other species: namely, low reproductive skew (Galezo et al., 2022). Thus, one question concerns why mountain gorillas do not appear to more strongly favor paternal siblings. Current evidence indicates that single-male gorilla groups across research sites are entirely genetically polygynous (reviewed in Rosenbaum & Silk, 2022), and while there can be considerable temporal variation, reproductive skew is generally much higher in multi-male gorilla groups than in, for example, chimpanzee, savannah baboon, or rhesus macaque groups (Vigilant et al. 2015, Surbeck et al. 2017, Alberts et al. 2003, Widdig et al. 2004). We propose that, despite possessing a mating system quite unlike these other primate species, mountain gorillas still exhibit a comparable maternal sibling bias due to a mismatch between their historical mating structure–which we speculate consisted of highly polygynous one-male units–and their contemporary social structure of tight-knit, often multi-male groups. In other words, while individuals in highly polygynandgrous groups might find it too difficult to detect and adjust affiliation toward paternal kin, perhaps mountain gorillas fail to do so because, until very recently, it was unnecessary. If co-residency was enough to identify patneral kin with reasonable accuracy, a more sophisticated recognition mechanism would be unlikely to evolve.

Notably, while we see little evidence that mountain gorillas show a prosocial bias towards paternal siblings, patterns of aggressive behavior suggest there may still be kin recognition mechanisms at play for all sibling types. Aggression remained low across most combinations of relatedness and sex configurations, with one exception: mixed-sex interactions among non-siblings. This pattern is consistent with males deploying aggression in the context of mate attraction or coercion. Past research in gorillas suggests male aggression towards females may have a number of non-mutually exclusive functions within those domains: to police female-female aggression, to discourage female dispersal or mate choice, or to indicate protective ability or overall condition (Robbins, 2009; Breuer et al., 2016). The fact that this kind of aggression was observed less frequently among related male-female pairs is another observation consistent with accurate kin discrimination. It also suggests active inbreeding avoidance, to the extent that aggression truly serves a mate attraction function. While death and dispersal have been suggested to obviate the need for sophisticated inbreeding avoidance mechanisms in some primates (e.g. baboons; Galezo et al., 2022), such an explanation is unlikely to apply to contemporary mountain gorillas. Living with opposite-sex relatives after sexual maturity is a routine occurrence in this species. Prior research confirms strong inbreeding avoidance between father-daughter dyads in this species (Vigilant et al. 2015), but further work is needed to investigate the extent to which male mate choice is manifested via female-directed aggression, and whether females, for their part, possess additional mechanisms to avoid mating with kin, including paternal siblings.

Together, these observations–prosocial biases towards kin that do not appear to be fully explained by familiarity; a stronger maternal than paternal sibling prosocial bias; and avoidance of intersexual aggression across all sibling types–both speak to key questions about the development of great ape sibling relationships and present two additional puzzles for interpretation. First, traditional mechanistic explanations for sibling biases that typically appeal to exposure during developmental periods appear largely inconsistent with our results and the nature of mountain gorilla sociality, in which siblings and non-siblings, and maternal and paternal siblings, are all likely to have significant exposure to one another during development. It is possible that early-life exposure effects via repeatedly sharing night nests (Fossey, 1979) function analogously to the manner in which co-residence duration serves as a key component of kin recognition in humans (Lieberman et al., 2007), or that preferential mother-father relationships post-birth might lead to social preferences among siblings (Rosenbaum et al., 2016). Individuals may also possess some degree of phenotype matching ability (Widdig, 2007; Parr et al., 2010; Langergraber, 2012; Pfefferle et al., 2014).

Second, the lack of evidence for a prosocial bias towards paternal siblings is not readily reconciled with clear behavioral evidence of reduced aggression within these same dyads. This remarkable disjunct between apparent sibling recognition and sibling bias suggests that from a mountain gorilla’s perspective, paternal siblings are known entities that nevertheless are less attractive social partners than maternal siblings, despite each being equal relatives. There may be multiple, non-mutually exclusive explanations for this dynamic. Perhaps the presence of paternal siblings provides fewer benefits to an individual than do other sibling types–this possibility, while previously suggested (e.g. Cords et al., 2018), has not been systematically investigated and is an ideal target for future research. A mismatch between historical and current social structure might also lead to inconsistent, weakened kin recognition among paternal siblings that manifests in the contrasting patterns we report. Ultimately, disentangling these potential explanations within a species that only exists in the wild may depend on the opportunity to study long-term mating patterns and the impacts of “natural experiments” such as early maternal loss or adoption (most often carried out by adult males in this species; Fossey, 1979, Morrison et al. 2021).

## Conclusion

Our analyses of sibling relationships in mountain gorillas provide extensive, large-scale information on the dynamics of cooperation and competition in a primate society where, as in humans, potential social partners vary greatly in the genes, developmental stage, and biological sex they share with each other. We find a selective sibling bias for prosocial behaviors, in that siblings who share matrilineal kinship affiliate at greater rates than either paternal half-siblings or non-siblings, and that this bias weakens as individuals become more distant in age. While such a result is consistent with a wide range of previous research, none of the reasons proposed for this selective bias in primates appear to apply to our population: mountain gorillas gain regular exposure to siblings of all types, across their entire lives; furthermore, patterns of aggressive behavior, in contrast to affiliation, suggest that mountain gorillas can in fact recognize paternal siblings, though they evidently do not favor them as cooperative partners. Ultimately, our study underscores a diversity of means, some evidently yet to be revealed, through which individuals might perceive and engage in sibling relationships to achieve fitness outcomes.

## Materials and Methods

Our study subjects came from a population of habituated wild mountain gorillas living in Volcanoes National Park, Rwanda, that have been monitored nearly continuously for the last 54 years by the Dian Fossey Gorilla Fund. Using focal follow and scan data collected by researchers and staff, we compiled a dataset of all available dyadic gorilla behavior spanning the years of 2003 to 2017. We then supplemented this dataset with demographic and relatedness data (for maternal relatedness, via direct observation; for paternal relatedness, via genetic paternity determination–see e.g. Vigilant et al., 2015) on individuals pulled from long-term records. From this combined dataset, we excluded interactions with infants <1 year of age at time of observation, parent-offspring interactions, and interactions between dyads for which we could not calculate relatedness from available data. This yielded a final, curated dataset containing 157 unique individuals (75F, 82M; average age at time of observation = 9.75 years) and 38,996 total hours of observation.

### Composition of dyads

Our dataset of behavior from 157 individuals contained 1957 unique dyad pairs. Of these dyads, 1258 shared neither a mother nor father (“non-siblings”), 555 shared a father but not a mother (“paternal siblings”), and 43 shared both a mother and a father (“full siblings”). In addition to dyads known to share a mother but not a father (n = 50), there were a number of dyads with the same mother, but with paternity data missing for one or both individuals (n = 51). To maximize sample size, we combined these two groups into the category of “maternal siblings”; due to this analytic choice, this category can be effectively conceived of as “at least maternal siblings”. See Tables S1 and S2 in Supplementary Materials for analyses using only confirmed maternal siblings, which were very similar to those reported below. Mixed-sex dyads were the most common sex category in our dataset (n = 981), followed by male-male (n = 520) and female-female (n = 456). Dyads differed in age by an average of 5.90 years (SD: 4.57 years; range: 0 – 23.5 years); for reference, the average interbirth interval in mountain gorillas is 3.9 years (Eckardt et al., 2016). We used this age difference variable as our primary index of familiarity between individuals. While we also had information on the natal groups of individuals, which could also serve as a potential index of familiarity, we do not focus on this variable in analyses, as it did not allow us to disambiguate between relatedness and familiarity–dyads of individuals who grew up in different natal groups were virtually never (n = 3) siblings in our dataset.

### Behavioral Measures

We evaluated five different categories of dyadic behaviors as outcome variables: grooming, playing, non-contact aggression, contact aggression, and time spent in close (2m) proximity. We operationalized these behaviors from standardized definitions used in previous publications about this gorilla population (see e.g. Rosenbaum et al., 2015). Trained observers regularly undergo interobserver reliability tests. The former four behavioral categories were evaluated as counts (corrected for exposure time; see *Data Analysis*) within the dyad during focal observations, regardless of directionality, while the latter category of time in close proximity was evaluated by counting the number of instantaneous scan samples in which a dyad was observed within 2 m of each other (also corrected for exposure time).

### Data Analysis

We conducted all analyses in R (version 4.1.2). Our main statistical models for each behavioral outcome consisted of cross-classified generalized linear mixed models (conducted using the *glmmTMB* package; Brooks et al., 2017) that included random intercepts for each individual within the dyad, as well as the dyad itself. Given low incidences of many behaviors, we aggregated behaviors into annual counts, making the dyad-year the fundamental unit of analysis (total n = 7858). Even with annual aggregation, instances of aggression were uncommon. Therefore, counts of contact and non-contact aggression were summed into a single category for analysis (see Tables S3 and S4 and Figures S1 and S2 in Supplementary Materials for results with individual aggression categories, which were qualitatively similar to those reported below).

In models predicting each behavioral outcome, we included terms for relatedness, age difference, and sex makeup, as well as two-way interactions between relatedness and sex makeup, relatedness and age difference, and sex makeup and age difference. As mothers plausibly mediate many of the social behaviors we examined, especially early in life, we also included the average proportion of observations with mothers in close proximity, and this variable’s interaction with relatedness, as covariates in all models. In models containing significant main effect or interaction terms, we decompose omnibus comparisons and report targeted marginal effects and contrasts using the *emmeans* package (Lenth, 2022), with all reported *p*-values corrected for false discovery rate. We modeled our count outcomes as rates with a negative binomial family in *glmmTMB* and offset term for exposure time (either logged hours of observation, or logged sum of scans for both individuals, per dyad-year). For each model, we verified model fit by inspecting the deviation, dispersion, and outliers of residuals using the *DHARMa* package (Hartig, 2022). All data and code necessary to reproduce our results are available publicly at https://osf.io/6qgj5.

## Acknowledgments

The authors are grateful to Winnie Eckardt and Robin Morrison for their expertise and assistance with data. We thank the Rwandan government and the Rwanda Development Board for their long-term support of the research, monitoring, and protection activities of The Dian Fossey Gorilla Fund’s Karisoke Research Center. We are deeply indebted to all Karisoke field staff for their tireless support in collecting long-term behavioral and demographic data. This study was supported by the University of Michigan, the NSF Graduate Research Fellowship Program and Doctoral Dissertation Improvement Grant No 1122321, and the donors who support The Dian Fossey Gorilla Fund.

## Competing Interests

The authors report no competing interests.

## Supplementary Analyses

Tables S1 – S2. Results for analyses using a ‘stricter’ categorization of maternal siblings (n = 50; see Methods for details of categorization; total dyad-years: 7625).

**Table S1.**
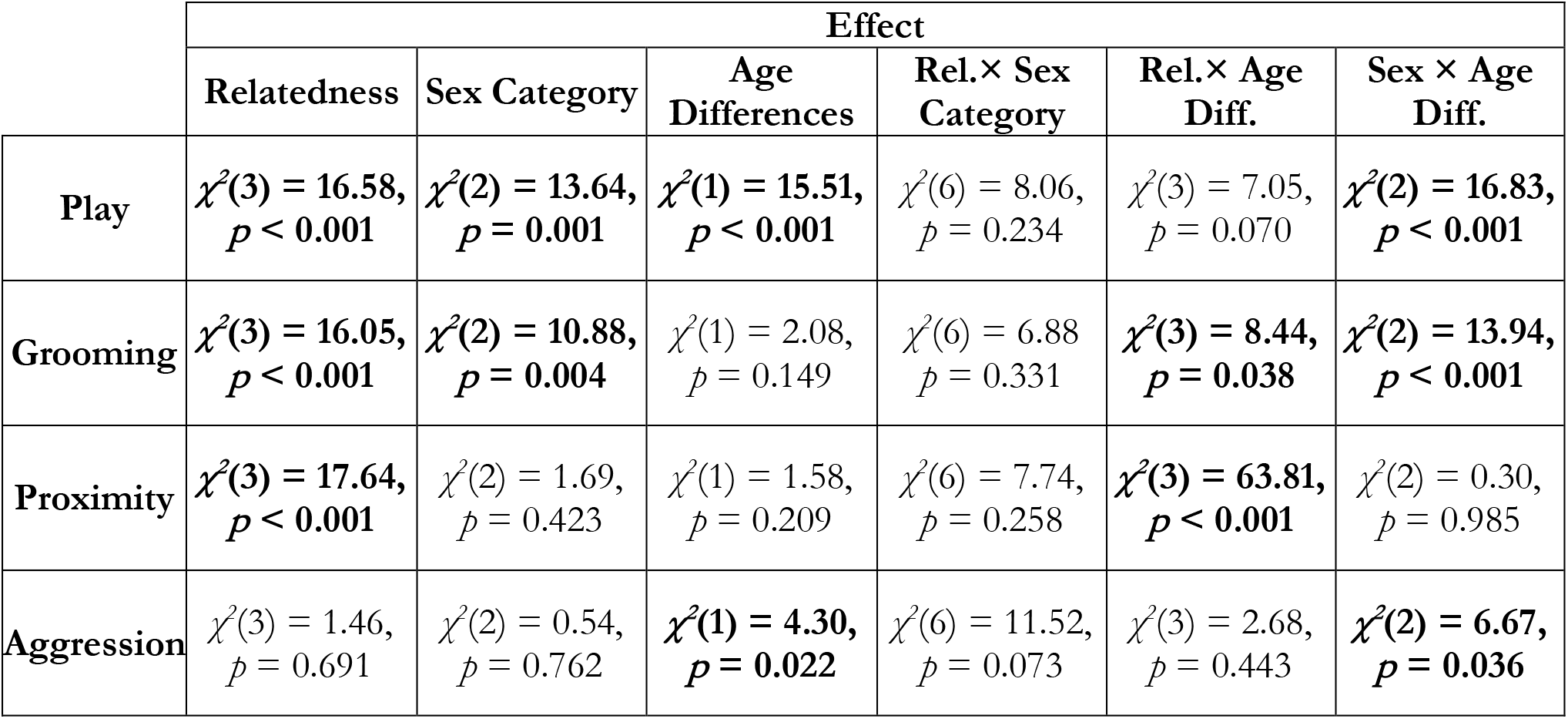
Omnibus statistics for target parameters (full model results available from data and code posted publicly at https://osf.io/6qgj5). Effects *p* < 0.05 bolded.

**Table S2.** Estimated marginal means and standard errors across relatedness and sex categories.

|  | Relatedness |  |  |  | Sex Category |  |  |
| --- | --- | --- | --- | --- | --- | --- | --- |
|  | Full Siblings | Maternal Half | Paternal Half | Non-Siblings | Female - Female | Male - Male | Mixed - Sex |
| **Play** | 2.91 (0.58) | 1.83 (0.42) | 1.42 (0.17) | 1.14 (0.11) | 1.17 (0.22) | 2.95 (0.50) | 1.46 (0.20) |
| **Grooming** | 2.51 (0.60) | 1.61 (0.45) | 0.30 (0.04) | 0.34 (0.04) | 1.49 (0.30) | 0.48 (0.10) | 0.73 (0.11) |
| **Proximity** | 44.9 (5.5) | 42.6 (5.9) | 15.8 (1.0) | 18.9 (1.0) | 30.2 (3.2) | 25.9 (2.6) | 26.4 (2.1) |
| **Aggression** | 0.43 (0.08) | 0.64 (0.13) | 0.56 (0.04) | 0.76 (0.05) | 0.50 (0.07) | 0.68 (0.09) | 0.60 (0.06) |

Tables S3 – S4, Figures S1–S2. Results for analyses assessing contact aggression and non-contact aggression separately (total dyad-years: 7822).

**Table S3.** Omnibus statistics for target parameters (full model results available from data and code posted publicly at https://osf.io/6qgj5).

|  | Effect |  |  |  |  |  |
| --- | --- | --- | --- | --- | --- | --- |
|  | Relatedness | Sex Category | Age Differences | Rel.× Sex Category | Rel.× Age Diff. | Sex × Age Diff. |
| **Contact Aggression** | $\chi^2(3) = 5.58, p = 0.134$ | $\chi^2(2) = 1.57, p = 0.456$ | $\chi^2(1) = 1.48, p = 0.223$ | $\chi^2(6) = 10.54, p = 0.104$ | $\chi^2(3) = 0.34, p = 0.953$ | $\chi^2(2) = 3.74, p = 0.154$ |
| **Non-Contact Aggression** | $\chi^2(3) = 2.96, p = 0.398$ | $\chi^2(2) = 3.35, p = 0.187$ | $\chi^2(1) = 1.92, p = 0.166$ | $\chi^2(6) = 10.23, p = 0.116$ | $\chi^2(3) = 1.17, p = 0.760$ | $\chi^2(2) = 2.22, p = 0.330$ |

**Table S4.** Estimated marginal means and standard errors across relatedness and sex categories.

|  | Relatedness |  |  |  | Sex Category |  |  |
| --- | --- | --- | --- | --- | --- | --- | --- |
|  | Full Siblings | Maternal Half | Paternal Half | Non-Siblings | Female - Female | Male - Male | Mixed - Sex |
| Contact Aggression | 0.23 (0.04) | 0.22 (0.04) | 0.28 (0.02) | 0.34 (0.02) | 0.22 (0.03) | 0.28 (0.04) | 0.30 (0.03) |
| Non-Contact Aggression | 0.10 (0.03) | 0.18 (0.04) | 0.18 (0.02) | 0.23 (0.02) | 0.18 (0.03) | 0.18 (0.03) | 0.13 (0.02) |

**Figure S1.**
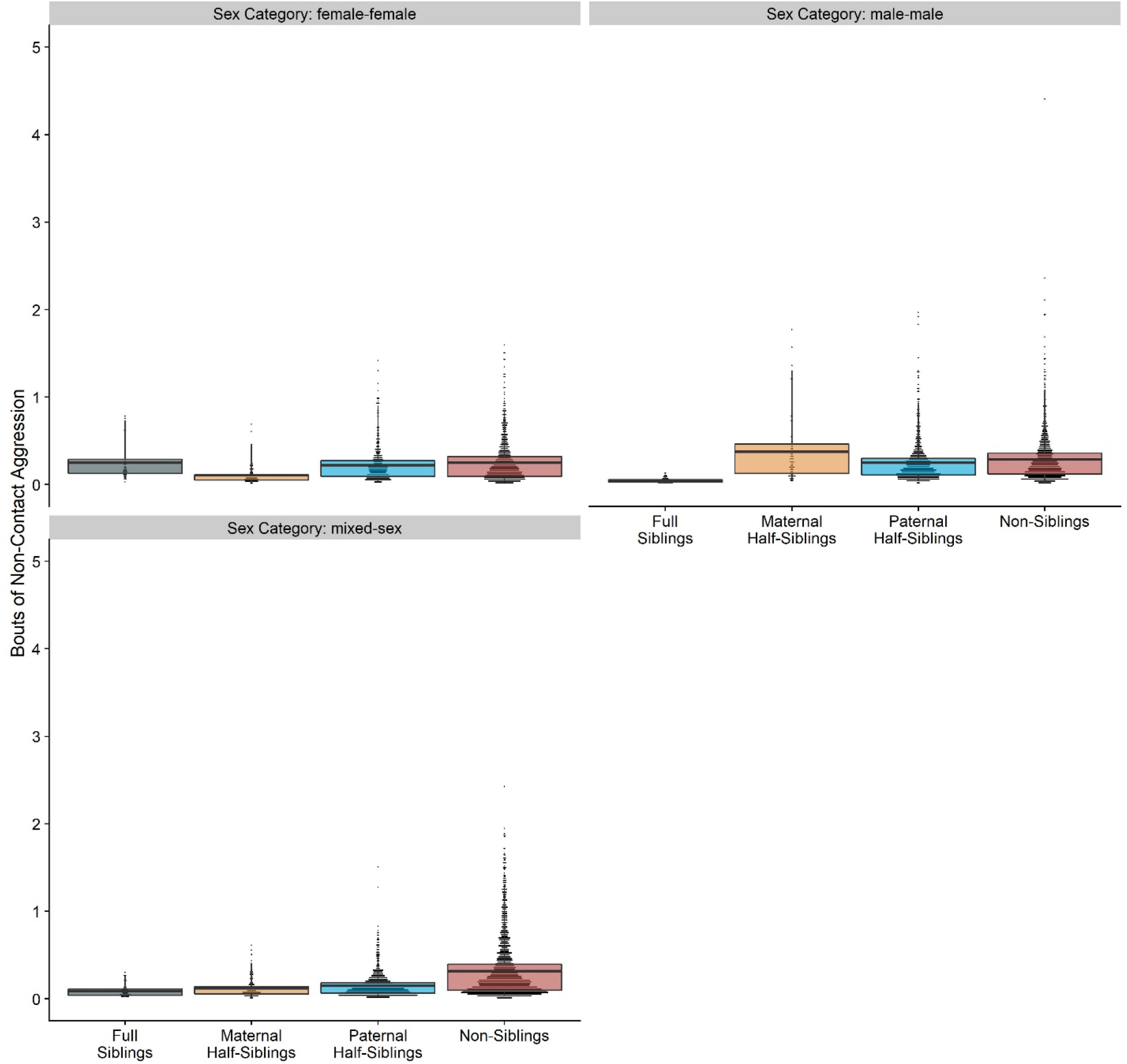
Box and dot plots showing estimated non-contact aggression within gorilla dyads, separated by relatedness and sex category.

**Figure S2.**
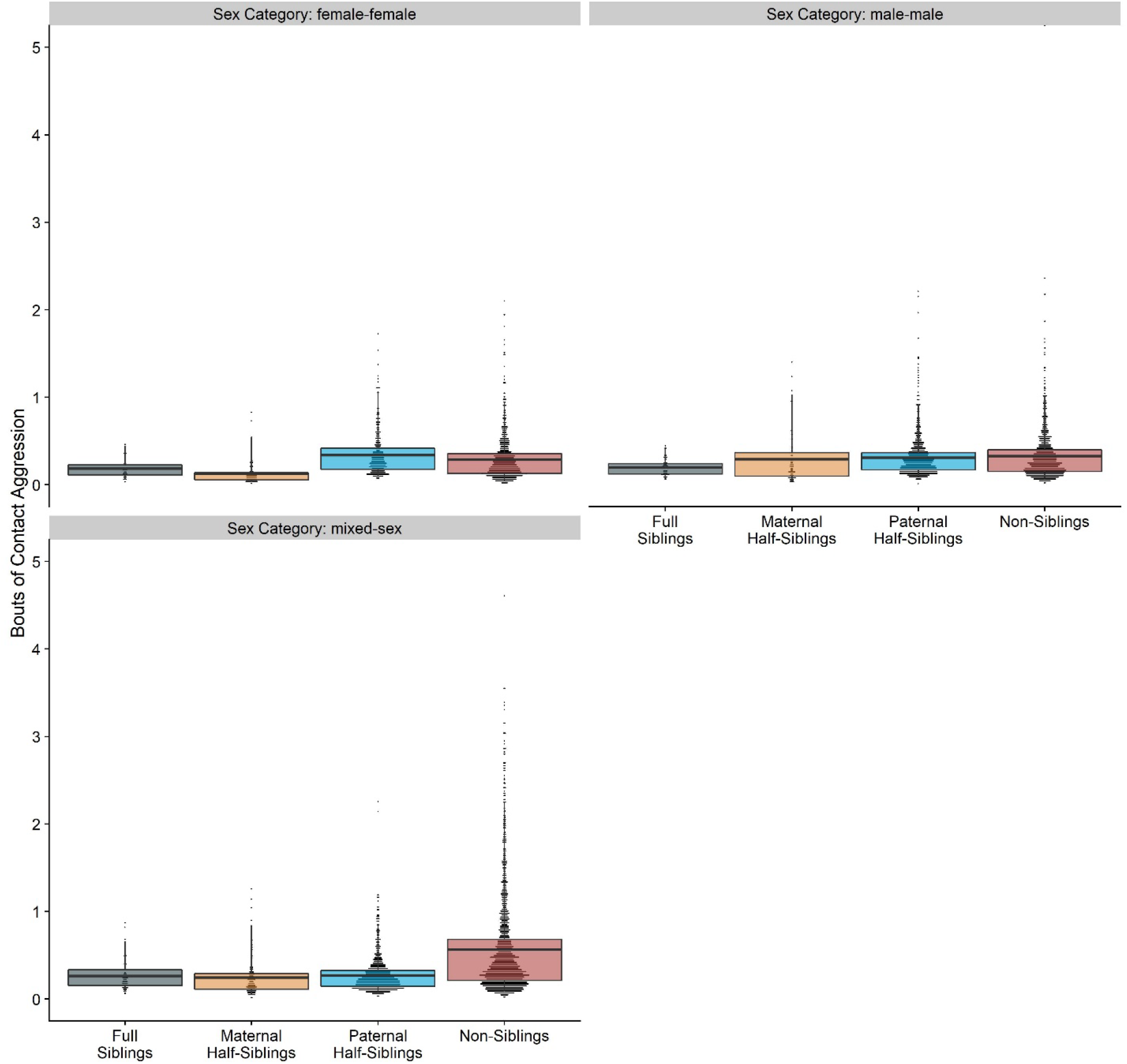
Box and dot plots showing estimated contact aggression within gorilla dyads, separated by relatedness and sex category.

## Notes

### Competing Interest Statement

The authors have declared no competing interest.

